# Toward Granular Brain Intrinsic Connectivity Networks and Insights into Schizophrenia

**DOI:** 10.1101/2025.06.11.659084

**Authors:** Shiva Mirzaeian, Kyle M. Jensen, Ram Ballem, Pablo Andrés Camazón, Jiayu Chen, Vince D. Calhoun, Armin Iraji

## Abstract

Spatial group independent component analysis (sgr-ICA) has become a crucial method to understand brain function in functional magnetic resonance imaging (fMRI) research, especially in resting-state fMRI (rsfMRI) studies. Early studies identified large-scale brain networks using sgr-ICA with lower order (e.g., 20 - 45 components); however, more recent studies have employed higher model orders (e.g., 200 components) to reveal more refined intrinsic connectivity networks (ICNs), offering a more detailed representation of functional brain architecture. This increased granularity has encouraged researchers to explore even higher model orders to better capture the brain’s function. Although previous studies explored higher model orders, small datasets often limited them. In this study, we addressed this gap by assessing an sgr-ICA model with 500 components using a large rsfMRI dataset of over 100,000 subjects. This extensive data set allowed us to provide a robust estimation of fine-grained ICNs. We further assessed diagnostic effects and cognitive performance using whole-brain functional network connectivity (FNC) of 502 individuals with schizophrenia and 640 typical controls from these ICNs. We also compared the results with ICNs obtained using a lower-order, multi-spatial-scale template. Results demonstrate that our approach yields a large set of reliable and fine-grained ICNs, enhancing characterization of schizophrenia related dysconnectivity patterns. Specifically, we observed a relatively large number of ICNs within the cerebellar and paralimbic area. We detected significant hypoconnectivity between the cerebellar and subcortical domains, including the basal ganglia and thalamic regions. We also found hyperconnectivity between the cerebellar domain and the visual, sensorimotor, and higher cognitive domains, as well as between the sensorimotor and subcortical domains. Our finding revealed that granular ICNs can detect significant FNC differences between cohorts which are missed in larger scale ICNs. This work highlights the capability of higher model order ICA to capture distinct, fine-grained ICNs, enriching our understanding of FNC and serving as a valuable addition to current multiscale ICN templates. The ICNs derived from this study may serve as valuable references for future research, with the potential to improve the clinical utility of rsfMRI and advance the study of psychiatric disorders.

## 1 Introduction

Models of brain communication in functional neuroimaging often rely on the assumption that temporal dependencies in neurophysiological data reflect functional connectivity [1]. Independent component analysis (ICA), a widely used multivariate method, plays a crucial role in uncovering the functional organization of the brain [2]. ICA is a blind source separation technique that decomposes data into maximally independent components [3]. Within neuroimaging research, it has proven valuable for identifying functional networks or intrinsic connectivity networks (ICNs) [2]. Two primary ICA-based approaches have been employed to estimate functional entities in the brain [4]. The first involves performing ICA on individual subjects followed by clustering to identify shared patterns across samples [5] [6]. While this approach accounts for individual variability, it is susceptible to challenges like inter-individual differences, variations in data acquisition, and the dynamic nature of brain function. The second, more robust method involves a group-level ICA framework, which creates shared functional entities across a cohort and subsequently back-reconstructs these patterns for individual subjects [7]. This approach has become a cornerstone for clinical and large-scale studies due to its reliability and consistency in capturing functional connectivity patterns across populations [4] [8] [9].

Earlier studies employing group-level ICA focused on low-order models (20 - 45 components) that identified large-scale networks, such as default mode and salience networks [10][2] [11]. However, the inherent complexity of brain networks suggests the presence of smaller, functionally distinct subnetworks embedded within these large-scale systems[12]; hence, Higher order ICA (75-200 components) has been employed to identify such fine-grained ICNs, providing a more detailed representation of the brain’s functional architecture [8] [13] [14] [15] [16]. Although some studies have explored even higher model orders (e.g., 500 or 1000 components), they are often applied to small datasets, highlighting the need for further extension to improve robustness and generalizability [17].

The advent of population-level neuroimaging has introduced massive datasets with terabytes of high-resolution brain images, enabling researchers to uncover the neural underpinnings of individual differences [4]. Large datasets improve the reliable estimation of ICNs and functional patterns by enhancing statistical power, reducing noise, and capturing more representative brain connectivity, which enables the derivation of robust, generalizable ICNs that better reflect individual and group differences. Such large-scale data requires efficient dimensionality reduction techniques to extract meaningful representations while managing computational challenges. Functional brain atlases, derived from these datasets, provide a structured framework to summarize complex connectivity patterns into image-derived phenotypes, which are critical for characterizing brain networks [18] [19]. Data-driven atlases from large datasets extracted from fMRI capture functional organization and individual variability and enhance reliability by improving statistical robustness and reducing noise [20]. Among these, high-dimensional ICA-based atlases effectively delineate spatially continuous and functionally specific regions, making them valuable for studying brain connectivity.

We recently analyzed a large dataset of over 100,000 subjects using multi-model order ICA, ranging from 25 to 200 components, to develop the multiscale NeuroMark functional atlas of the brain [4]. Subsequently, the functional atlas was refined, organized, and labeled, with each network described using terminology familiar to cognitive and affective neuroscience, resulting in NeuroMark-fMRI-2.2 [20]. Building upon this foundation, we extend the ICA framework by implementing a group-level ICA at model order 500 on the same dataset to extract a replicable and reliable set of fine-grained ICNs. This extension results in a new template named NeuroMark-fMRI-500, which offers enhanced granularity and provides additional functional insights beyond those captured in NeuroMark 2.2.

### 1.1 Schizophrenia

Schizophrenia is a complex psychiatric disorder that impacts thinking, emotions, and daily functioning [21]. It is diagnosed based on a combination of symptoms, as there is no single test for the condition. The primary symptoms include positive symptoms such as hallucinations, delusions, and disorganized speech or behavior, and negative symptoms like lack of motivation, blunted affect, and emotional withdrawal, along with significant social difficulties [21] [22] [23]. Schizophrenia profoundly affects brain function by disrupting normal network connectivity, leading to patterns of hypo- and hyper-connectivity and reducing the brain’s ability to integrate and process information efficiently [24] [25]. For example, dysconnectivity between the motor cortices and cerebellar areas is a typical feature observed in schizophrenia [26]. Additionally, dysfunction in the triple network, which includes the central executive network, the default mode network, and the salience network, has been implicated in the disorder, with their interactions often being deficient in schizophrenia [27] [28] [8]. Although diverse methods, such as graph theory [29], decomposition techniques, and seed-based analyses [30], have been used to study abnormal functional integration across brain circuits in schizophrenia, sgr-ICA has played a crucial role in identifying and characterizing schizophrenia-related patterns of functional connectivity. For instance, multiscale ICA, which investigates functional sources across multiple spatial scales, has uncovered sex-specific differences in schizophrenia [16]. Similarly, Telescopic ICA, a novel method employing a recursive ICA strategy that leveraged information from larger ICNs to guide the extraction of smaller ICNs, revealed significant associations in the posterior cortex and precuneus regions linked to auditory hallucinations in individuals with schizophrenia [8]. Using higher orders in ICA, which involves analyzing a large number of fine-grained components, allows for the identification of more subtle and complex brain network interactions, shedding light on changes in functional connectivity associated with schizophrenia [4][17].

In this study, we analyze functional network connectivity (FNC) from 1,142 subjects, including typical controls (TC) and individuals with schizophrenia (SZ), extracted using NeuroMark-500 and its association with cognitive scores. Our results show that NeuroMark-500 captures a relatively large number of ICNs within the cerebellar area, revealing their strong hypoconnectivity with subcortical-extended thalamic regions as well as subcortical-basal ganglia regions, and hyperconnectivity with the sensorimotor cortex. These findings underscore the cerebellum’s significant role in brain function and its disruption in schizophrenia.

## 2 Method

### 2.1 Data collection and data preparation

For this study, we relied on the same dataset and quality control (QC) procedures as described in [4], without introducing any modifications. Resting-state fMRI (rsfMRI) datasets were used from 100,517 subjects, sourced from over twenty private and public datasets. A complete list of these datasets, along with details on accessing additional information, is provided in Supplementary Section 1 in [4]. These datasets originate from cohorts with varying sex and diagnosis ratios, age distributions, and imaging protocols that differ in spatial and temporal resolution. The QC criteria included (a) a minimum of 120 time points (volumes) in the rsfMRI time series, (b) mean framewise displacement less than 0.25 mm, (c) head motion transitions within 3º rotation and 3 mm translation in any direction, (d) high-quality registration to an echo-planar imaging template, and (e) spatial overlap between individual masks and the group mask, including the top and bottom ten slices, exceeding 80 %. These criteria were chosen for their feasibility across diverse datasets [4]. Applying these established QC criteria resulted in 57,709 individuals (57.4 %) passing the QC requirements, forming the QC-passed dataset. Data pre-processing follows the procedures described in [4] [31]. When available, preprocessed data from a given dataset were used; otherwise, preprocessing pipelines were applied. The preprocessing steps included rigid body motion correction, slice timing correction, and distortion correction, using the FMRIB Software Library (FSL v6.0, https://fsl.fmrib.ox.ac.uk/fsl/fslwiki/) and the Statistical Parametric Mapping (SPM12, https://www.fil.ion.ucl.ac.uk/spm/) toolboxes within the MATLAB environment. Next, preprocessed subject data were warped into the Montreal Neurological Institute (MNI) space using an echo-planar imaging (EPI) template. This approach has been shown to outperform structural templates [32] when distortion correction is unavailable or unfeasible, which was the case for this study. Finally, subject data were resampled to 3 mm^3^ isotropic voxels and spatially smoothed using a Gaussian kernel with a 6 mm full width at half-maximum (FWHM).

### 2.2 Independent component analysis

We performed spatial Group-level ICA (sgr-ICA) using the Group ICA Toolbox (GIFT) [33] [34] on data. The steps for sgr-ICA are as follows. First, we applied variance normalization on voxel time courses and conducted subject spatial principal components analysis (PCA) to retain the principal components (PCs) with subject-level variance exceeding 95 %. Next, we performed group spatial PCA by concatenating subject PCs to further reduce the dimensionality of the data and decrease the computational demands of sgr-ICA [35]. We used a memory-efficient subsampled time PCA (STP) approach due to data size to calculate Group PCs [36]. Next, we ran sgr-ICA using the Infomax algorithm [37] with model order 500, resulting in 500 fine scale components.

### 2.3 Creating the ICN Template

We used 500 independent components to identify the set of replicable fine-grained ICNs as NeuroMark-500 template. First, we assessed the reliability and quality of the components using the ICASSO index quality (IQ) [38]. Components with IQ value below 0.80 were also excluded from further analysis. Secondly, we evaluated the stability of the components using a split-half approach.

We randomly split the QC-passed data into two independent halves and applied sgr-ICA separately on each half, resulting in 500 components from each half. Next, we identified the best-matching components between two independent half splits by assessing spatial similarity using Pearson correlation with full sample as reference. This process was repeated 50 times. Components with an average similarity below 0.80 [39] across the 50 iterations from the full sample were excluded. By applying these two criteria— IQ thresholding and split-half ICA stability—we obtained a subset of components that were replicable and reliable. Finally, to classify components as ICNs, we applied the same procedure in [31][4], selecting components that exhibited high spatial overlap with gray matter and low spatial similarity to motion artifacts, ventricular signals, or other known artifact patterns. Additionally, for components located in cortical regions, we ensured that their peak activation to be within gray matter area. Based on these, 131 ICNs were identified as NeuroMark-fMRI-order-500 template. The identified ICNs are categorized into domain and subdomain labels according to NeuroMark 2.2 template [20] using spatial similarity measured by Pearson correlation.

After establishing these replicable ICNs, we leveraged them to study their application in FNC and group comparisons between TC and SZ.

### 2.4 Clinical application

We utilized the BSNIP dataset [40] [41] for further analysis, which comprises 1,142 subjects, including TC and SZ. Participants underwent the Structured Clinical Interview for the Diagnostic and Statistical Manual of Mental Disorders IV (DSM-IV), and assessments were conducted while participants were clinically stable. To estimate subject-specific independent components and time courses, we employed multivariate-objective optimization ICA with reference (MOO-ICAR), leveraging the NeuroMark-500 components as spatial priors. This approach has demonstrated effectiveness in capturing subject-specific information while reducing artifacts [42]. We then applied time course cleaning procedures, including detrending and despiking, to eliminate drifts, sudden fluctuations, and residual artifacts [43] [44]. Finally, we computed static FNC (sFNC) matrices by calculating pairwise correlations between the cleaned time courses of different brain domains across the entire dataset [45] [15].

To statistically assess group differences in sFNC between TC and SZ, first, we applied a linear regression model to regress out potential confounding factors, including age, sex, scanning site, and mean framewise displacement. Next, we conducted two-sample t-tests on the cleaned sFNC to compare the TC and SZ groups. Additionally, we evaluated the relationships between cleaned sFNC and cognitive performance measures using Pearson correlation coefficient. We removed group effects by demeaned both cleaned sFNC and cognitive scores to ensure that the observed associations reflect relationship between variables rather than being skewed by group-level variations. Cognitive performance was measured using standardized z-scores from the Brief Assessment of Cognition in Schizophrenia (BACS) [46].This analysis enabled us to explore how functional connectivity differences relate to distinct cognitive domains.

### 2.5 ICN comparison: NeuroMark-500 vs. NeuroMark 2.2

We compared ICNs from the NeuroMark-500 with ICNs from NeuroMark 2.2. NeuroMark 2.2 is a reliable and replicable multi-spatial-scale ICNs template using sgr-ICA with order ranging from 25 to 200. We performed comparisons based on spatial map size, component distribution across functional domains, and the ability to capture group differences. ICN size was defined by the number of voxels exhibiting z-scored intensity values greater than 1.96 (Z *>*1.96). To evaluate the ability to capture group differences, we extracted subject-specific time courses of NeuroMark 2.2 ICNs from the same dataset used in our group comparison analysis and applied an identical methodological pipeline to both approaches, ensuring a consistent and direct comparison.

## 3 Results

### 3.1 ICN results

Based on the criteria outlined in Section 2.3, we identified 131 ICNs extracted from NeuroMark-500. Figure 1a presents the spatial maps of these ICNs, organized into domains and subdomains according to the NeuroMark 2.2 template. Among these, 27 ICNs were classified within the cerebellar domain. The higher cognitive and paralimbic domains also exhibited substantial contributions, with 23 and 22 ICNs, respectively.

**Figure 1:**
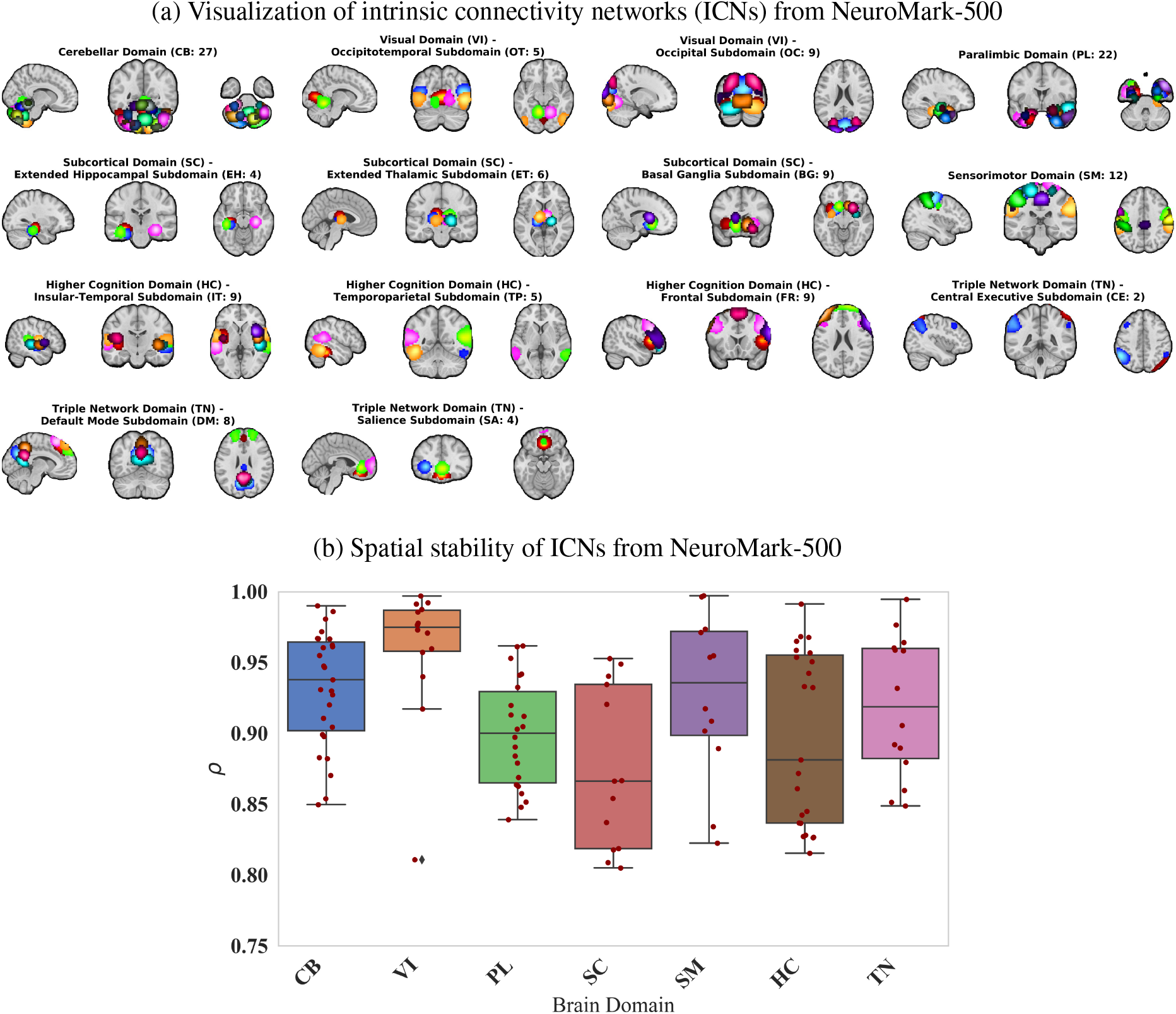
(a) Representation of 131 spatial maps of ICNs from NeuroMark-500 for each of the seven domains and 14 subdomains-labeling based on based on NeuroMark 2.2 template: cerebellar (CB), visual-occipitotemporal (VI-OT), visual-occipital (VI-OC), paralimbic (PL), subcortical extended hippocampal (SC-EH), subcortical-extended thalamic (SC-ET), subcortical-basal ganglia (SC-BG), sensorimotor (SM), higher cognition-insular temporal (TC-IT), higher cognition-temporoparietal (HC-TP), higher cognition-frontal (HC-FR), triple network-central executive (TN-CE), triple network-default mode (TN-DM), and triple network-salience (TN-SA). The spatial maps of ICs are thresholded with a z-score of 1.96 (p-value *<* 0.05). (b) Spatial stability of the 131 ICNs from NeuroMark-500, evaluated across 50 split-half iterations.

Figure 1b presents the stability index, which quantifies the spatial similarity of each ICN generated from 2 half split across 50 iterations. This metric provides insight into the consistency of ICN identification and reflects the robustness of the analysis. The results indicate that the stability index for all ICNs exceeds 0.80, demonstrating high reproducibility. ICNs within the visual domain exhibited the highest stability (*>*0.95), followed by those in the cerebellar, sensorimotor, and triple-network domains, which also showed strong stability.

### 3.2 Clinical application results

Figure 2 presents the sFNC matrix and the group comparison analysis between cohorts using the NeuroMark-500 template from the BSNIP sample. The sFNC heatmap displays pairwise correlations between ICN time courses, providing a detailed view of functional connectivity patterns across all subjects. The accompanying group comparison plot illustrates the cleaned sFNC differences between TC and SZ, based on the statistical approach described in2.4. The group comparison analysis reveals both hyperconnectivity and hypoconnectivity patterns in the SZ group relative to TC. Notably, there is pronounced hypoconnectivity between the cerebellar domain and both the subcortical-extended thalamic and subcortical-basal ganglia regions. Conversely, the SZ group exhibits significant hyperconnectivity between the cerebellar domain and sensorimotor, insular-temporal, and temporoparietal subdomains. Additionally, the thalamic region shows significant increased connectivity with sensorimotor, insular-temporal, and temporoparietal areas, highlighting widespread alterations in cross-subdomain functional integration.

**Figure 2:**
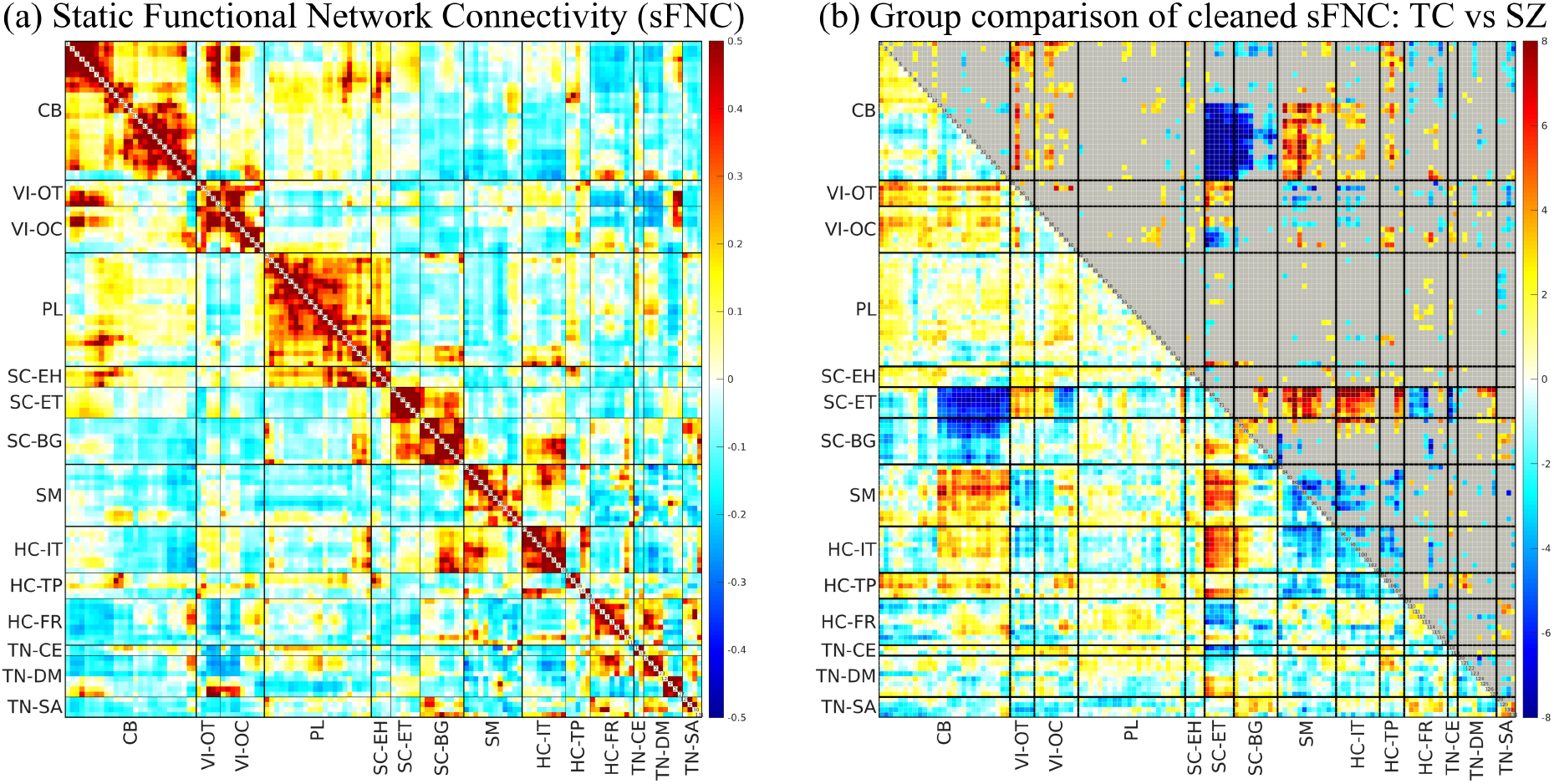
Clinical application results of ICNs from NeuroMark-500 for 1,142 subjects in the BSNIP dataset. (a) Heatmap of sFNC between ICNs, showing averaged pairwise correlations between time courses of different brain domains. (b) Group comparison of cleaned sFNC (TC vs. SZ) using NeuroMark-500: The lower triangle represents *−* log(p-value) × sign(t-value), while the upper triangle highlights connectivity with p-values *<* 0.05. P-values are FDR corrected. Blue indicates hypoconnectivity in the SZ, while red represents hyperconnectivity in the SZ. Labels: cerebellar (CB), visual-occipitotemporal (VI-OT), visual-occipital (VI-OC), paralimbic (PL), subcortical extended hippocampal (SC-EH), subcortical-extended thalamic (SC-ET), subcortical-basal ganglia (SC-BG), sensorimotor (SM), higher cognition-insular temporal (HC-IT), higher cognition-temporoparietal (HC-TP), higher cognition-frontal (HC-FR), triple network-central executive (TN-CE), triple network-default mode (TN-DM), and triple network-salience (TN-SA).

Figure 3 illustrates the correlations between cleaned sFNC and cognitive performance after regressing out diagnostic effects from both variables. The connectograms display highly significant Pearson correlations (p *<* 0.005), with p-values corrected for multiple comparisons using the false discovery rate (FDR) method. Cognitive measures include overall cognition, verbal memory, working memory, and processing speed. The correlation analysis shows a generally consistent pattern for all cognitive associations. The plots exhibit a significant positive association between sFNC in subcortical-extended hippocampal and subcortical-basal ganglia with cerebellar domain, indicating that strong connectivity in these regions supports cognitive function. Conversely, cerebellar-sensorimotor, thalamic-sensorimotor and thalamic-insular temporal, and thalamic-frontal is negatively correlated with the same cognitive measures, suggesting that increasing connectivity between these networks may be linked to cognitive impairments.

**Figure 3:**
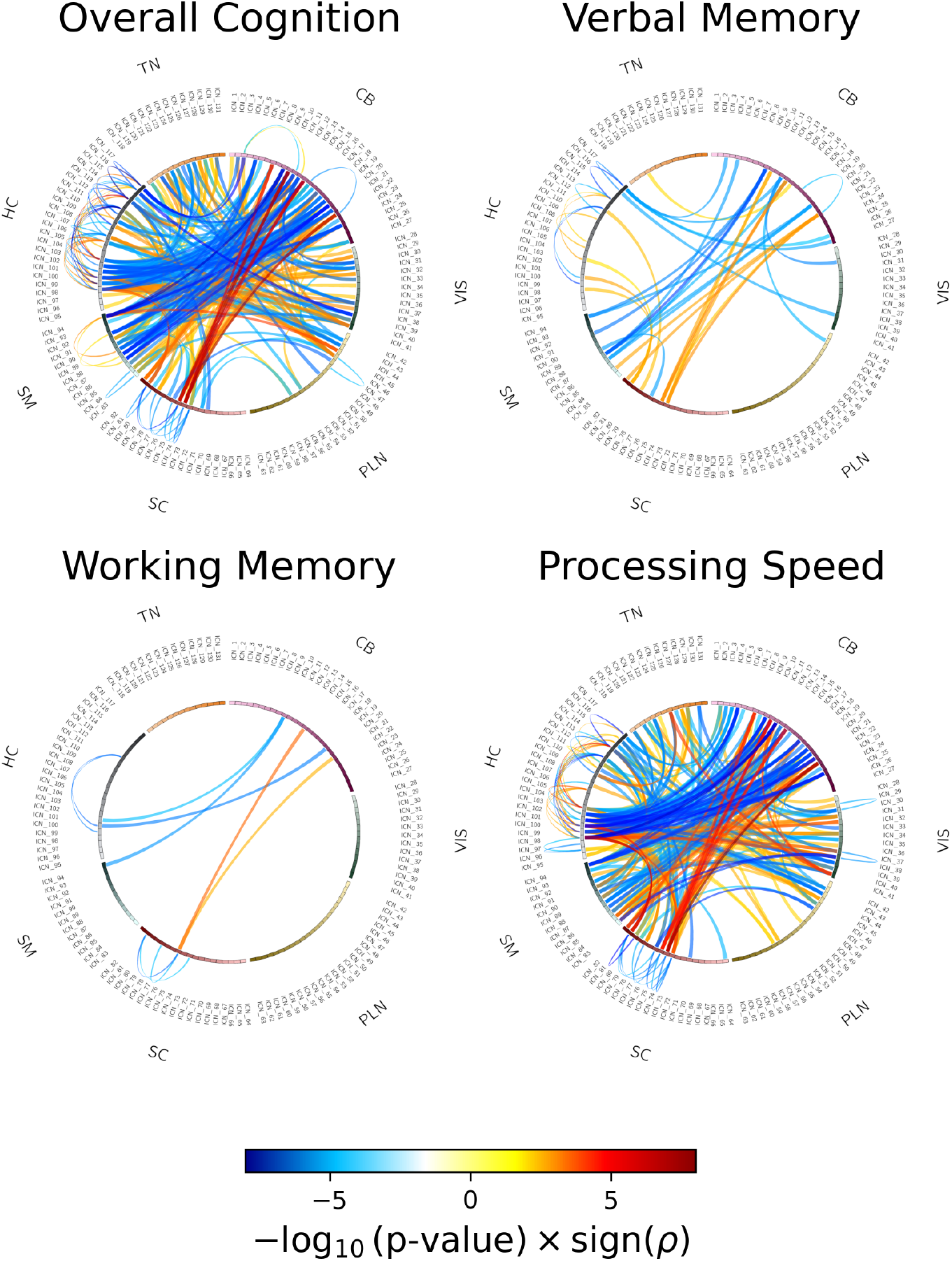
Associations between cognitive scores and cleaned sFNC from NeuroMark-500.The connectograms highlight Pearson correlation coefficients between cleaned sFNC and cognitive measures—including overall cognition, verbal memory, working memory, and processing speed—showing significant correlations (p *<* 0.005). p-values are FDR corrected.

### 3.3 Comparison of ICNs from NeuroMark-500 vs. NeuroMark 2.2

Figure 4a compares the ICNs generated using the NeuroMark-500 with ICNs produced by NeuroMark 2.2, focusing on the number of ICNs identified within each brain domain and ICN size. The analysis reveals that ICNs derived from the NeuroMark-500 are in general smaller in size across all brain domains. Additionally, NeuroMark-500 approach identifies a greater number of ICNs across cerebellar, visual, paralimbic, subcortical and higher cognitive domain. This combination of smaller ICN sizes and an increased number of ICNs highlights the NeuroMark-500 capacity to achieve finer granularity in brain network parcellation.

**Figure 4:**
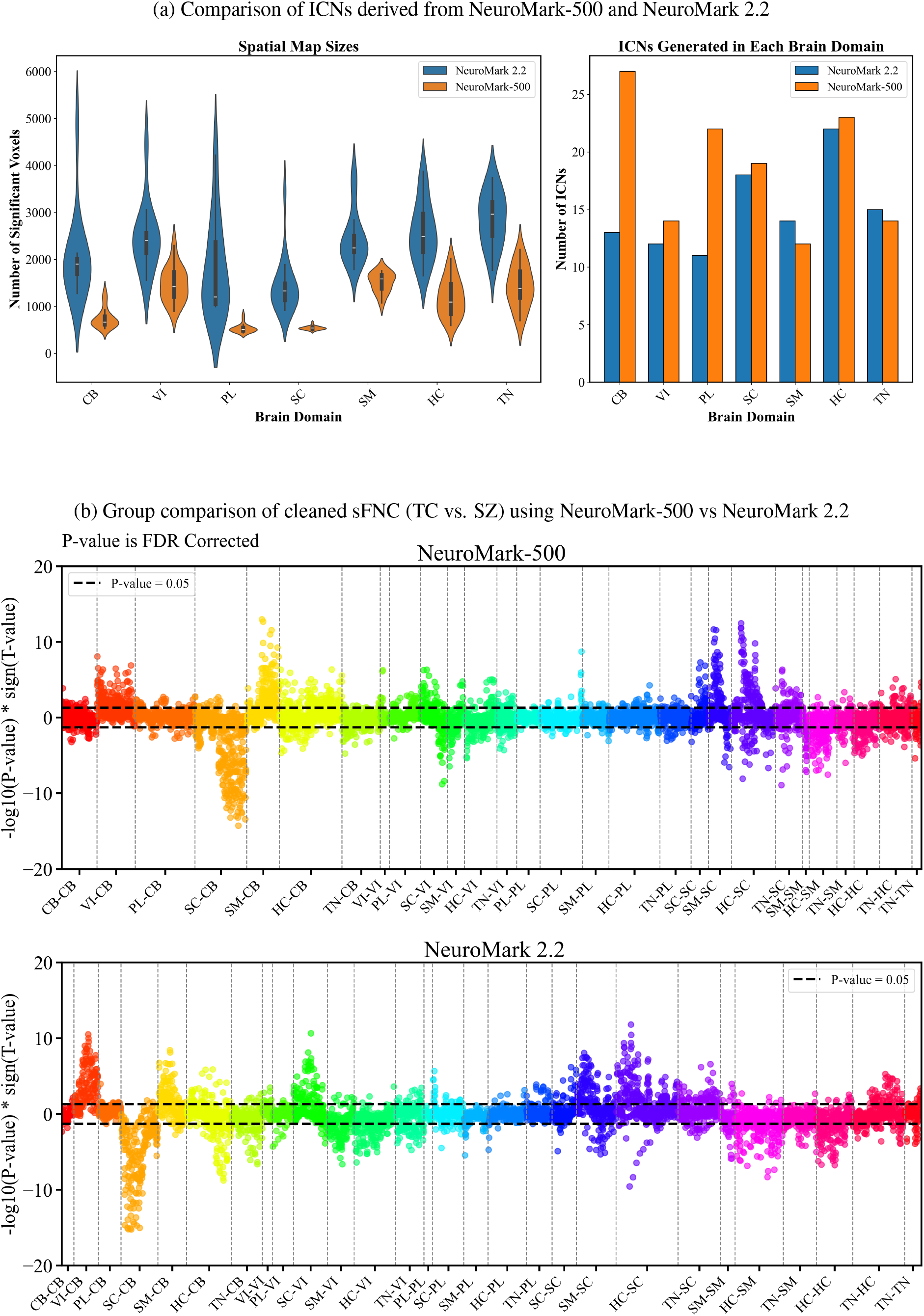
Comparison of ICNs derived from NeuroMark-500 and NeuroMark 2.2. (a) Spatial map sizes and ICNs distribution across brain domains. (b) Group comparison of sFNC (TC vs. SZ) using NeuroMark-500 vs NeuroMark 2.2: Positive side indicates hypoconnectivity in the SZ, while negative side represents hyperconnectivity in the SZ.

Figure 4b compares the group comparison results between TC and SZ from NeuroMark-500 vs NeuroMark 2.2. The plot shows that the two approaches exhibit a generally consistent pattern, which provides additional support for the validity and interpretability of the results obtained using NeuroMark-500 framework. However, notable differences are observed in the paralimbic and cerebellar domains, where the results extracted from NeuroMark-500 reveals stronger and more widespread diagnostic effects—particularly in connections between the cerebellar and sensorimotor domains, and between the paralimbic and cerebellar domains. The results also indicate that three brain domains—cerebellar, sub-cortical, and sensorimotor—demonstrate diagnostic associations with multiple other brain domains. Specifically, the cerebellar domain shows significant hypoconnectivity with the subcortical domain, higher cognitive networks, and the triple network. Additionally, a significant hypoconnectivity is observed between the higher cognitive and sensorimotor domains. On the other hand, significant hyperconnectivity involving the cerebellar domain is observed with the visual, sensorimotor, and higher cognitive domains. The sensorimotor domain also exhibits significant hyperconnectivity with the sub-cortical domain.

Figure 5 presents two representative examples highlighting the capability of NeuroMark-500 to capture finer-grained brain networks and provide more detailed insights into the functional patterns of the SZ group, compared to the larger scale ICNs used in NeuroMark 2.2. In each example, we selected one ICN from NeuroMark 2.2 and identified its correspondence into two distinct ICNs from NeuroMark-500 using Pearson correlation of spatial maps.

**Figure 5:**
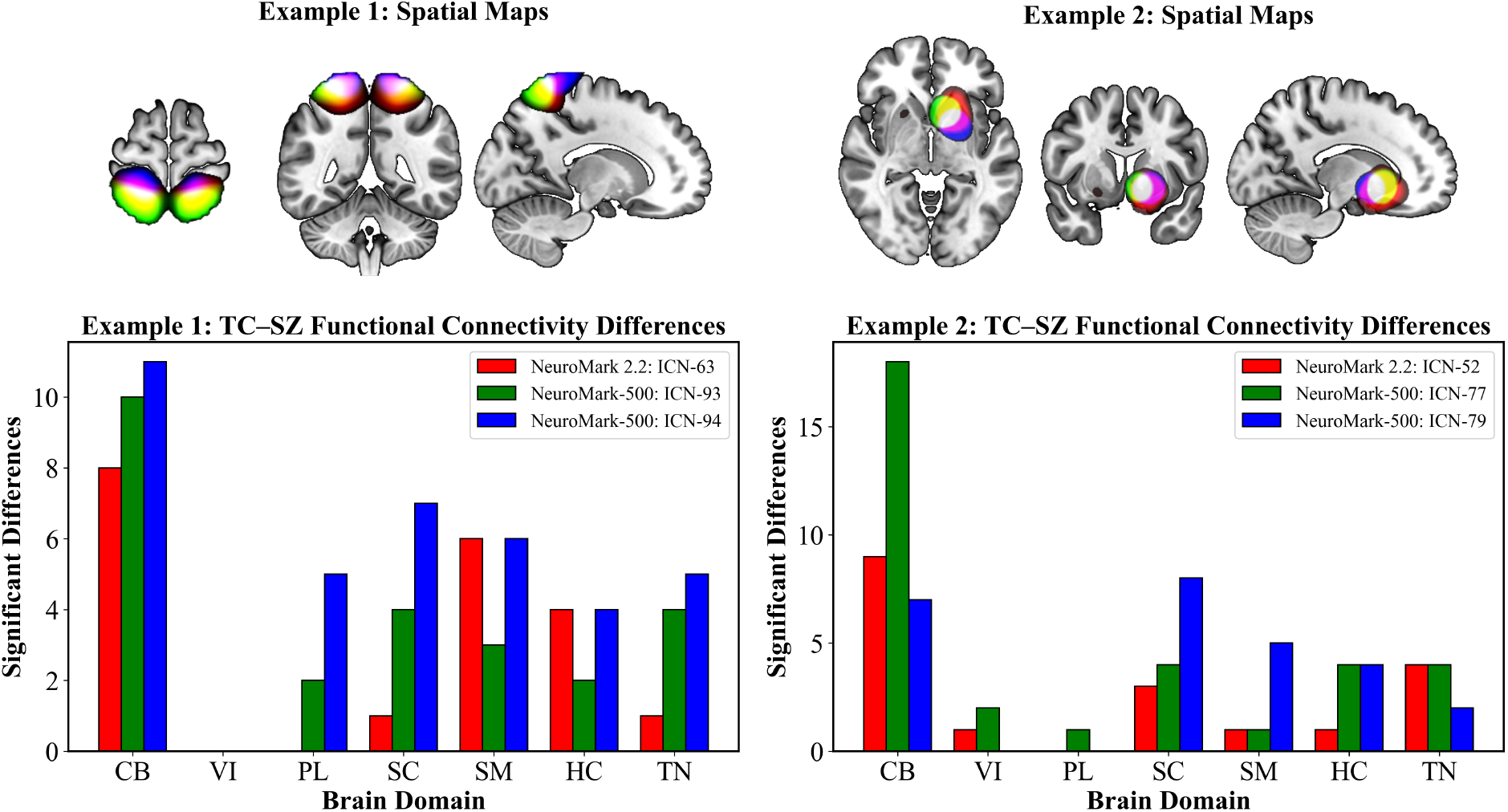
Barplot illustrates the number of significant functional network connections showing group differences between TC and SZ from the group comparison analysis. Each example compares an ICN from NeuroMark 2.2 with its two most spatially similar ICNs derived from NeuroMark-500. The NeuroMark-500 decomposition reveals finer-grained ICNs and identifies additional significant group differences highlighting its enhanced sensitivity for capturing clinically relevant connectivity alterations. Red represents spatial maps from NeuroMark 2.2, blue and green show spatial maps from NeuroMark-500, yellow is overlap between red and green, magenta is overlap between red and blue, white is overlap between all three.

In Example 1, the sensorimotor ICN (ICN 63) from NeuroMark 2.2 showed spatial correlations of 0.73 and 0.55 with two sensorimotor ICNs (ICN 93 and ICN 94, respectively) from NeuroMark-500. In Example 2, the subcortical–basal ganglia ICN (ICN 52) from NeuroMark 2.2 exhibited the highest spatial similarity with ICNs 77 and 79 from NeuroMark-500, with correlation coefficients of 0.61 and 0.59, respectively. These spatial maps visually and quantitatively confirm that increasing the ICA model order allows a single ICN to branch out into multiple more localized components, thereby revealing more detailed functional brain organization. We also examined diagnosis association between cohorts across the corresponding ICNs. The NeuroMark-500 revealed additional significant diagnostic associations that were not captured by NeuroMark 2.2. In both examples, ICNs derived from NeuroMark-500 capture significant diagnostic effects within the paralimbic domain that are not detected using the NeuroMark 2.2 template. Additionally, the number of connectivity pairs showing significant group differences is higher when using the finer-grained ICNs from NeuroMark-500 compared to the coarser, large-scale ICNs, highlighting the benefits of increased spatial resolution in detecting subtle group effects.

## 4 Discussion

This study demonstrates the efficacy of higher order ICA for resting-state fMRI data, identifying 131 ICNs with high granularity. The high reproducibility of ICNs, as evidenced by a stability index exceeding 0.80 across 50 runs, underscores the robustness of this method. Our findings highlight the prominent involvement of the cerebellum, with 27 out of 131 networks spatially overlapping with this region. Although, historically understudied in neuroimaging research, emerging studies have established the cerebellum as a major hub in the brain’s functional circuitry [47] [48] [49], more recently, subregions in the cerebellum have been established as having specialized roles in integrating and processing information [47] [48] [49]. The relatively large number of ICNs we identified within the cerebellum underscores its significant role in brain function and lends further support towards future research aiming to explore the functional contributions of the cerebellum in cognition.

Beyond the cerebellum, our results also reveal a notable enrichment of ICNs within the paralimbic domain, with 22 ICNs classified under this category. The paralimbic domain, centered around regions adjacent to limbic structures—including the entorhinal cortex, medial temporal lobe, and temporal pole—is known to support a wide range of higher-order cognitive functions such as emotion regulation, memory, language, and learning [50] [51]. The substantial number of ICNs identified within the paralimbic domain emphasizes its critical role in overall brain function and further motivates future investigations into its specific contributions to cognition. Crucially, our group comparison analysis in figure 5 revealed significant differences between TC and SZ cohort specifically within the paralimbic and subcortical domains—differences that were not detected using the NeuroMark 2.2 template. This finding highlights the added value of higher order ICA, which enables the extraction of more focal ICNs, thereby increasing sensitivity to subtle alterations in functional connectivity.

Additionally, our comparison of sFNC between TC and SZ using NeuroMark 2.2 and NeuroMark-500 as more fine scale ICNs reveals that, while the overall pattern remains similar, a key distinction arises in the hippocampus subdomain. The NeuroMark-500 appears to enhance the separation of the hippocampus from neighboring regions. In contrast, the larger ICNs may capture effects influenced by adjacent structures, particularly the thalamus. This suggests that increased granularity may offer a more precise delineation of functional networks, which could be critical for accurately characterizing the role of the hippocampus in psychopathology-related connectivity alterations.

Moreover, our results reveal strong hypoconnectivity within subcortical regions and the cerebellar, along with hyperconnectivity in sensorimotor-cerebellar areas. These connectivity disruptions align with prior studies reporting similar dysconnectivity patterns in schizophrenia [52] [53] [54], reinforcing the critical role of subcortical and cerebellar dysfunction in the disorder. This underscores the need for more targeted investigations into these regions and their contributions to schizophrenia-related network alterations.

Our findings further suggest that schizophrenia-related cognitive deficits are closely linked to altered subcortical and cerebellar connectivity. The observed hypoconnectivity within thalamic, basal ganglia, and cerebellar may indicate impaired integration of these key subcortical structures with the rest of the brain, potentially disrupting executive function, attention, and motor coordination. Given that these regions play a vital role in cognitive control and memory, their reduced connectivity may contribute to widespread cognitive impairments observed in schizophrenia [55]. Moreover, the positive correlation between subcortical-cerebellar connectivity and cognitive scores suggests that stronger functional interactions between these areas benefit cognitive performance. This finding aligns with previous research emphasizing the role of the thalamus and basal ganglia in cognitive regulation [56, 57]. Conversely, hyperconnectivity between CB and SM networks was negatively correlated with cognitive function, which may reflect a compensatory but inefficient mechanism [55]. Excessive sensorimotor-cerebellar engagement may interfere with higher cognitive functions rather than enhance them, mirroring prior findings linking disrupted CB-SM interactions to cognitive deficits in schizophrenia [48]. Overall, these results reinforce the idea that schizophrenia is characterized by widespread dysconnectivity across subcortical, cerebellar, and sensorimotor networks. Distinct patterns of hypo- and hyperconnectivity appear to play a crucial role in cognitive dysfunction, highlighting the complex interplay between brain network disruptions and schizophrenia-related cognitive deficits.

This study opens several avenues for future research. First, it is important to note that NeuroMark-500 is based on a single, high model order decomposition, which allows for the extraction of a large number of fine-grained ICNs. On the other hand, the NeuroMark 2.2 template utilizes a multi-model-order ICA strategy, combining components derived from different lower model orders. There is significant promise in developing a unified network template that merges them Such a combined template could capitalize on the strengths of both approaches—capturing both large-scale, robust networks and subtle, focal ICNs—offering a more comprehensive and flexible tool for mapping brain functional architecture in both typical and clinical populations. Second, future work should explore even higher model orders to further refine network granularity, provided that sufficiently large datasets and rigorous validation procedures are employed to ensure robustness and reproducibility. Finally, the observed modular organization within the sFNC matrix involving the cerebellar domain, suggests the possible existence of subdomains within this region. Future research should explore this structure in greater depth and aim to classify the cerebellar domain into distinct, functionally meaningful subdomains.

## 5 Conclusion

This study demonstrates the power of fine-grained ICNs from large resting-state fMRI datasets in understanding brain functions. By applying a 500-component ICA model to an extensive dataset of over 58,000 individuals, we achieved 131 highly reproducible and spatially refined ICNs of the brain and introduced NeuroMark-500. Furthermore, our analysis of sFNC in schizophrenia revealed distinct patterns of hypo- and hyperconnectivity, particularly within subcortical, cerebellar, and sensorimotor regions, highlighting their critical role in schizophrenia-related cognitive deficits. These findings suggest that NeuroMark-500 can serve as a valuable template for future neuroimaging studies, with potential clinical applications in psychiatric research.

## 6 Ending sections

### 6.1 Data and Code Availability

For this study, we relied on the same dataset as described in [4], without introducing any modifications. A complete list of these datasets, along with details on accessing additional information, is provided in Supplementary Section 1 in [4].

### 6.2 Author Contributions

Shiva Mirzaeian: Conceptualization, Methodology, Investigation, Visualization, Writing – original draft. Kyle M. Jensen: Writing – review and editing, Visualization. Ram Ballem: Writing – review and editing, Visualization. Pablo Andrés Camazón: Writing – review and editing. Jiayu Chen: Writing – review and editing. Vince D. Calhoun: Conceptualization, Investigation, Supervision, Writing – review and editing. Armin Iraji: Conceptualization, Investigation, Supervision, Writing – review and editing.

### 6.3 Funding

This work was supported by the National Institutes of Health (NIH) under Grants R01MH136665 and R01MH123610, and by the National Science Foundation (NSF) under Grant No. 2112455.

